# A Robust Crystallographic Platform for High-Throughput β-Catenin Ligand Discovery

**DOI:** 10.1101/2025.07.13.663865

**Authors:** Marta Klejnot, Alicja N Skowron, Jaromir Szymański, Filip Ulatowski, Michał Biśta, Sylvain Cottens, Gabriela Dąbrowiecka, Daria Gajewska, Karolina M Górecka-Minakowska, Daria Kotlarek, Kinga Leszkowicz, Martyna W Pastok, Magdalena Sypień, Igor H Wierzbicki, Janusz Wiśniewski, Michał J Walczak

**Affiliations:** Captor Therapeutics S.A., ul. Duńska 1,54-427 Wrocław, Poland

**Keywords:** TDP – Targeted Protein Degradation, SBDD – Structure-Based Drug Design, crystallographic fragment screening, β-catenin, drug discovery, x-ray crystallography

## Abstract

This study presents a robust crystallographic platform for assessing compounds binding to β-catenin. We developed a standardized protein production protocol for the armadillo domain of β-catenin (BC-ARM) and performed biophysical screens using Surface Plasmon Resonance (SPR) and Differential Scanning Fluorimetry (DSF). These findings led to the successful determination of the co-crystal structure of BC-ARM with compound 1 binding to previously reported site but distinct from known transcription factor binding sites. To broaden the search for novel BC binding sites, we utilized FragLites library with a cyclic peptide-stabilized BC-ARM construct. This yielded two high-resolution co-crystal structures identifying a previously unreported binding hotspot. Recognizing the limitations of the cyclic peptide-bound construct for general screening, we designed a novel, truncated BC-ARM construct. This new construct eliminates unstructured regions, reliably producing high-quality, diffracting crystals suitable for high-throughput crystallographic studies. In conclusion, the ligand-bound β-catenin structures and this novel, robust BC-ARM construct establish a powerful platform for further β-catenin investigation.

## Introduction

Canonical Wnt/β-catenin pathway is a key regulator during embryonic development cell, differentiation but also in tissues homeostasis (1). β-catenin (beta-catenin, BC) has a double function, on one side it is involved in cell adhesion, on the other it is responsible for gene transcription. As part of the Wnt/β-catenin pathway it acts as a central signalling molecule linking Wnt mediated pathway to target gene transcription in the nucleus. Upon nuclear localisation, β-catenin interacts with transcription factors, like TCF/LEF family (T-cell factor/lymphoid enhancer factor) to induce Wnt target gene expression by recruitment of transcriptional co-activators (2), namely CBP and BCL9 (3,4). In normal, healthy cells β-catenin is tightly regulated through degradation by tumor suppressor protein APC (Adenomatous Polyposis Coli). APC is a scaffolding protein forming a destruction complex together with Axin, Casein Kinase I (CK1) and Glycogen Synthase Kinase 3β (GSK-3β) (5). β-catenin is phosphorylated by Casein Kinase I (CK1) and GSK-3β leading to β-catenin ubiquitination and subsequent proteasomal degradation, preventing its accumulation in the cytoplasm and nucleus and activation of target gene transcription (6). Aberrant canonical Wnt signaling causes accumulation of β-catenin causing it to become a key oncogenic driver for certain cancers, for instance in ∼80% colorectal cancers are somatic mutations in APC (7–9). As a results of loss-of-function mutation in APC, impaired recruitment of β-catenin to the destruction complex is observed and β-catenin stabilizes and accumulates in nucleus.

β-catenin is encoded by CTNNB1 gene. It is circa 85 kDa protein and consists of 12 armadillo (ARM) repeats, each consisting of ∼42 amino acid (10). All repeats form a characteristic superhelical structure, which generate a positively charged grove critical for binding partners such as TCF/LEF transcription factors, E-cadherin (11,12) and APC **(Figure 1)**. In addition to armadillo repeats β-catenin contains N- and C-terminal domains, which are flanking the ARM domain. The termini regions modulate β-catenin stability and activity. Post-translational modifications, like phosphorylation in these regions influence β-catenin activation or degradation. For instance, phosphorylation at SER552 by AMPK promotes stabilization of the protein by enhancing TCF/LEF-mediated transcription, whereas phosphorylation on SER33 and SER37 by HIPK2 and GSK3B triggers for proteasomal degradation (13). Protein Data Bank (PDB) consists multiple entries of β-catenin structures with various binding partners providing valuable insights into its molecular interactions and function. The crystal structure of β-catenin bound to TCF4 transcription factor (PDB ID: 2GL7) revealed how gene transcription in the Wnt signalling pathway is regulated. Structural insights helped to design small molecule targeting the β-catenin/TCF4 complex followed by its destabilization and degradation to treat some cancers, for instance molecule LF3 (a 4-thioureido-benzenesulfonamide derivative) reduced tumor growth and induced differentiation in a mouse xenograft model of colon cancer (14). However, besides being a useful tool to study Wnt/β-catenin signaling, there is a limited clinical information about further developments of this inhibitor. Furthermore, structural studies of APC binding to phosphorylated form of β-catenin (PDB ID: 1TH1) revealed how β-catenin is promoted to degradation via Ubiquitin-Proteasome System (UPS). Additionally, cell adhesion role of β-catenin and its cell-cell adhesion and tissue integrity was understood by crystal structure of β-catenin to E-cadherin (PDB ID: 1I7X), linking it to the actin cytoskeleton at adherens junctions (1). Relatively recently Kessler *et al.* (*15*) reported a crystal structure of β-catenin bound to compound 6 (PDB ID: 7AFW), which was a result of their screening campaign. The binding site of this compound was located between armadillo repeats two and three, near the BCL9 and TCF4 binding sites and was implicated as a promising perspective towards heterobifunctional degraders development (**Figure 1**).

**Figure 1.**
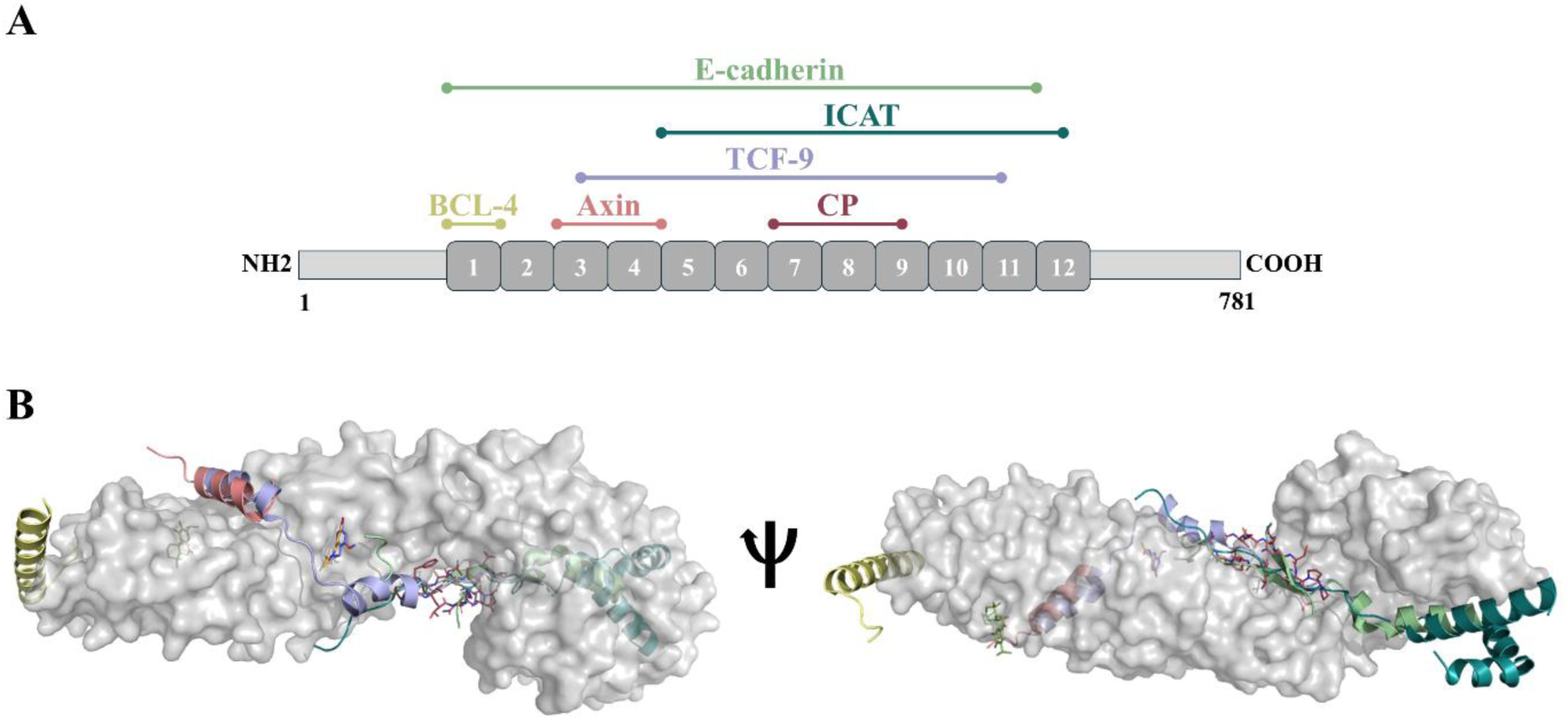
Interaction partners of β-catenin in relation to the binding sites of compounds 1, 2 and 3. *(A)* Schematic representation of binding partners of β-catenin. *(B)* Selected interactors in relation to crystal structures reported in this report - β-catenin (134-671) shown as gray surface, cyclic peptide (CP) shown in raspberry sticks. Other binding partners are shown in cartoon: Axin (salmon), BCL9 (pale yellow), E-cadherin (green), ICAT (teal) and TCF-4 (blue). Binding sites of the compounds are marked in boxes: compound 1 (olive sticks), compound 2 (orange sticks) and compound 3 (violet sticks).

Due to its role in Wnt signaling and their cancer driven implications, β-catenin is an interesting drug discovery target. Therefore, various β-catenin targeting options have been explored quite broadly to date. Even though multiple efforts and resources has been put into discovery of small molecules targeting β-catenin, none of the campaign was truly successful and β-catenin by being a challenging target has been deemed as undruggable. The primary shortcoming is that β-catenin has a flat protein surface and lacks well defined pockets that makes it difficult to target by a traditional drug discovery approaches. Its surface is largely positively charged hence there is a potential for a relatively high rate of false positives. Additionally, β-catenin is not only phosphorylated at multiple sites (14) but also due to its role in signal transduction and adhesion, it shares an extensive interaction surface for many protein binding partners, function which cannot be also disrupted by small molecules (1,15).

Heterobifunctional protein degraders and molecular glues are a new class of agents that eliminate, rather than just inhibit, their target proteins. In general, protein degraders mediate a transient interaction between an E3 ligase and a target protein leading to ubiquitination of this target protein and its subsequent degradation by proteasome (UPS) (16). Molecular glues are monovalent small molecules that induce the degradation of target protein, whereas heterofunctional protein degraders, are bivalent. They share overall mechanism of action. First protein degraders entered clinical testing in 2019, and the clinical trial landscape is rapidly evolving with several promising candidates already in clinical stages for various cancers (17,19). Moreover, they have been powerful tool for understanding biology (18). Targeted protein degradation also presents a promising approach for β-catenin degradation. Here we report development of a new crystallographic platform for drug discovery purposes of β-catenin. We determined a crystal structure of apo β-catenin using a new construct. Additionally, we report three crystal structures with small molecules. Noteworthy one of compounds binds in the same binding site as compound 6 reported by Kessler *et al.* (PDB:7AFW) **(Figure 1)** (15). These compounds could be a good starting point for development of heterobifunctional β-catenin degraders.

## Results and discussion

This study describes the successful establishment of a robust crystallographic platform for the reproducible assessment of beta-catenin-(BC)-binding compounds facilitating their advancement toward targeted protein degradation (TPD). The methodology involved the initial selection of specific compound series and the concurrent development of a standardized protein production protocol applicable to both biophysical and crystallization studies. A comprehensive literature review guided the selection of initial compounds which subsequently informed the design and synthesis of a series of analogs with systematic modifications to their side chains.

We employed carnosic acid, a previously reported beta-catenin binder (32), compound 6 from (15), and compounds from the FragLites library (33). Initial biophysical characterization of compounds identified in literature and their analog library was performed using Surface Plasmon Resonance (SPR) and Differential Scanning Fluorimetry (DSF) using armadillo domain BC-ARM (134–671). First, we performed the DSF screen at a constant fragment concentration of 500 μM or compound concentration of 100 μM that unfortunately revealed no hits. None of the compounds showed a real shift of denaturation temperature, which would be an indicator of binding and consequent protein stabilization (see **Table 1** for compounds selection). In parallel, SPR was used as an orthogonal screening method. In performed experiments we observed a response in dose-dependent manner; however, we struggled with a linear SPR response to a few compounds, followed by a weak signal and weak beta-catenin binding. Wherever possible, we determined the binding constant by the steady-state affinity binding model. Dose-response was observed also for some FragLites in SPR experiments, where we tested 19 fragments from which 2 compounds were identified as active with pKD∼3.5; 8 compounds showed dose-response, however the 1:1 fit was not possible; compound 3 showed irregular response; compound 2: KD = 241E-06 [M], Rmax = 17.2 (theoretical Rmax = 25RU) and compound 7: KD = 80E-06 [M], Rmax = 15.7, theoretical Rmax = 25 RU. Results of compound 6 agreed with previously published results by Kessler *et al*. (15).

**Table 1.**
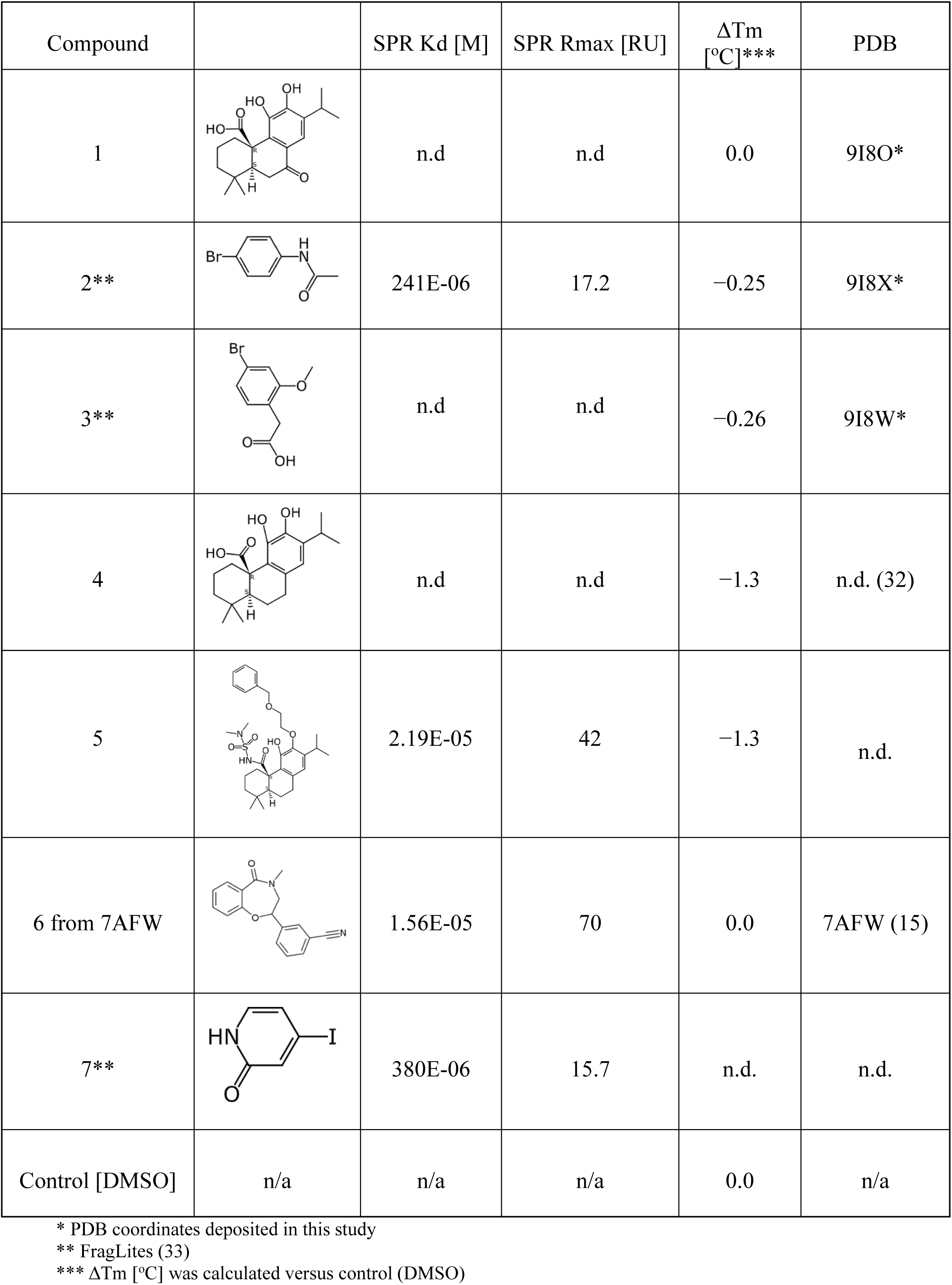
Chemical structure, affinity data and DSF data for compounds 1-7.

| Compound | | SPR Kd [M] | SPR Rmax [RU] | $\Delta T_m$<br>[°C]*** | PDB |
| --- | --- | --- | --- | --- | --- |
| 1 |  | n.d | n.d | 0.0 | 9I8O* |
| 2** |  | 241E-06 | 17.2 | -0.25 | 9I8X* |
| 3** |  | n.d | n.d | -0.26 | 9I8W* |
| 4 |  | n.d | n.d | -1.3 | n.d. (32) |
| 5 |  | 2.19E-05 | 42 | -1.3 | n.d. |
| 6 from 7AFW |  | 1.56E-05 | 70 | 0.0 | 7AFW (15) |
| 7** |  | 380E-06 | 15.7 | n.d. | n.d. |
| Control [DMSO] | n/a | n/a | n/a | 0.0 | n/a |

\* PDB coordinates deposited in this study
\*\* FragLites (33)
\*\*\* $\Delta T_m$ [°C] was calculated versus control (DMSO)

These preliminary findings informed subsequent crystallization experiments. Broad screening of commercially available crystallization conditions, combined with various protein-ligand mixtures and standard crystallization techniques, led to the successful determination of the co-crystal structure of β-catenin (141-305; R4) with compound 1 at 2.65 Å resolution (**Figure 2)**. Structural analysis revealed that compound 1 binds within a loop region situated between armadillo repeats 2 (residues 205–208) and 3 (residues 242–251), a binding mode consistent with previously reported β-catenin ligand interactions (*e.g.*, PDB ID: 7AFW). We identified the prominent hydrogen bond between the carboxyl group of the compound 1 and the hydroxyl group of SER246. Further stabilization of the position of the ligand was provided by a weaker CH-π interaction involving the 3-isopropylcatechol ring and the side chain of PRO247. These interactions orient the dimethyl cyclohexane ring of compound 1 towards the target, near the thioether side chain of MET243 and the main chains of THR205 and ASN206. In addition to the key interaction with SER246, the carboxyl group is situated within a hydrophobic pocket formed by VAL208, VAL248, and VAL251, while the 3-isopropylcatechol ring is solvent-exposed, suggesting a potential site on the isobutane group for chemical modification. Importantly, this binding site of compound 1 does not overlap with the recognition sites of transcription factors (TCF4 and BCL9), known to bind β-catenin, which is significant for therapeutic development as it does not interfere with those interactions (**Figure 1).**

**Figure 2.**
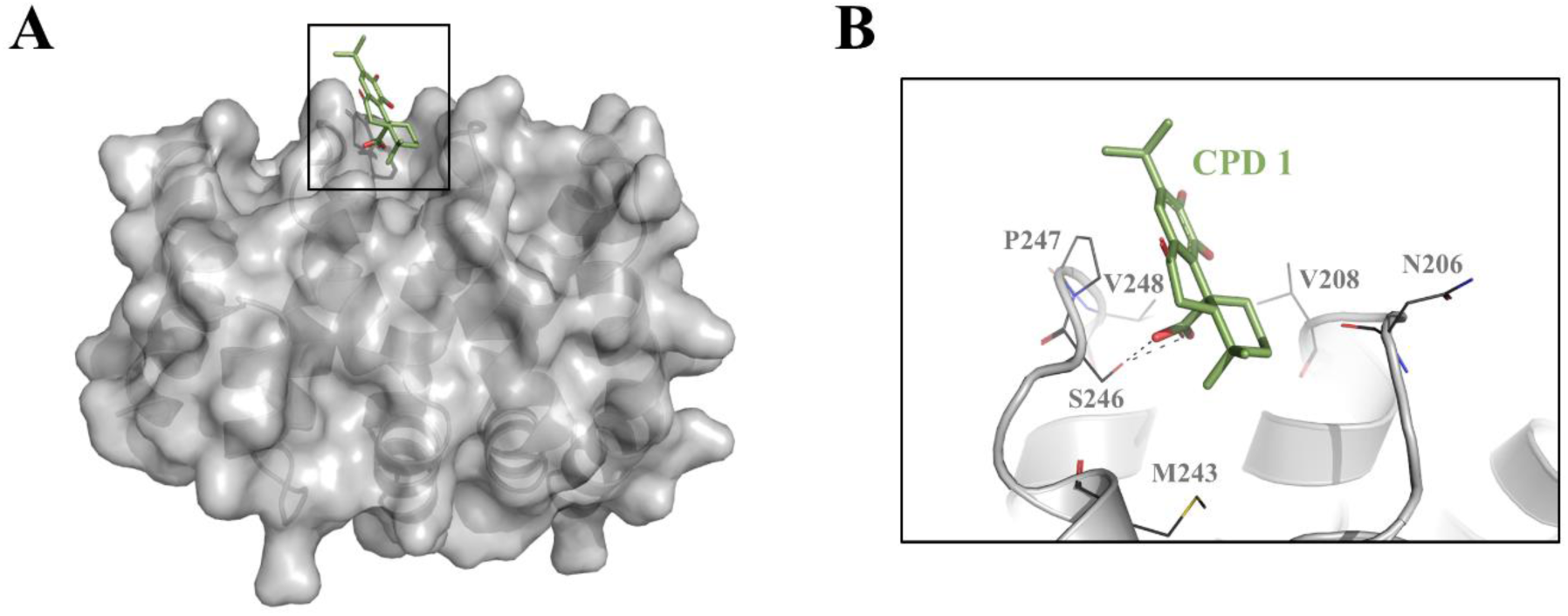
Overview of the crystal structure of β-catenin (141-305) bound to compound 1. *(A)* Representation of complex - β-catenin (141-305) shown as gray cartoon with surface bound to compound 1 shown in olive sticks. *(B)* Detailed representation of interactions between β-catenin (141-305) (gray cartoon) and compound 1 (olive sticks). Interacting amino acids are indicated by gray lines; bonds are indicated by black dashes (PDB: 9I8O).

To broaden the scope of β-catenin surface search and identify potential novel binding sites, we have decided to utilize a fragment-based screening, using the FragLites technology (33). FragLites is a library of small, halogen-containing fragments that can point to previously unidentified binding sites and guide its further development. To maximize a chance of obtaining high-resolution complex structures and allowing its confident modeling we have decided to use co-crystallization of the beta-catenin armadillo domain (BC-ARM, residues 141-305) bound to stabilizing macrocyclic peptide inhibitor (PDB ID: 7AR4). Using this system, we were able to obtained two high-resolution structures of BC-ARM in complex with compound 2 and compound 3 (**Table 2**: ARM-CP-Compound 2 – 1.98 Å, ARM-CP-Compound 3 – 1.89 Å) both occupying the same binding spot, in close proximity to ARG342, therefore identifying a β-catenin binding hotspot, which was not previously reported (**Figure 3**). The fragment oxygen from carbonyl/amide group is hydrogen-binding to GLU302, while the bromine ion lays near LYS345, ARG376 and ASN380 offering possible growing vectors to expand compound, gaining additional interactions and affinity.

**Figure 3.**
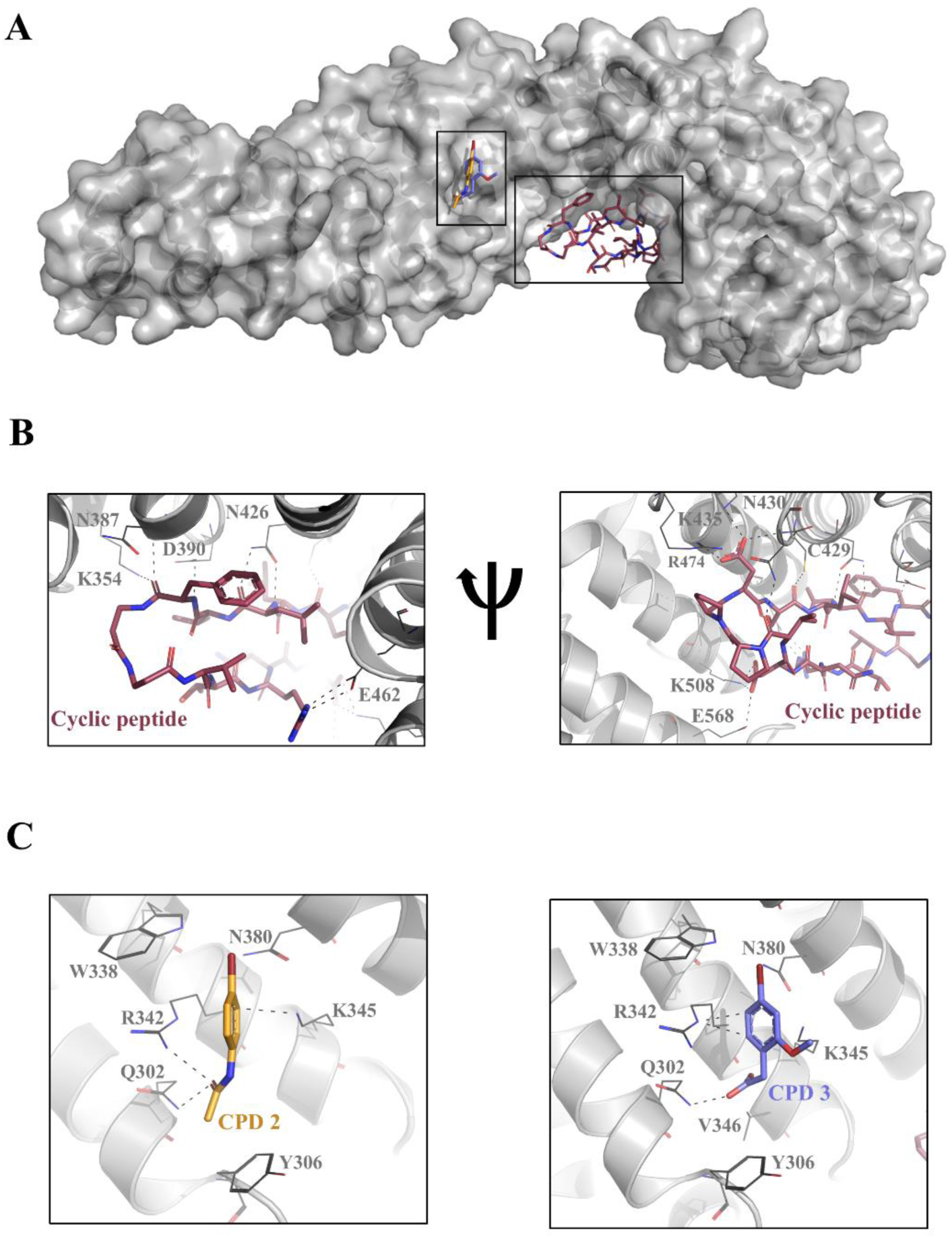
Overall crystal structure of beta-catenin (134-671) in a complex with cyclic peptide bound to compound 2 or compound 3. *(A)* Representation of full complex - beta-catenin (134-671) shown as gray cartoon with surface in complex with cyclic peptide (CP) shown in raspberry sticks bound to compound 2 (orange sticks) or compound 3 (violet sticks). *(B)* Detailed representation of beta-catenin (134-671) interactions with cyclic peptide. beta-catenin (134-671) amino acids involved in interactions with the cyclic peptide are indicated by gray lines; bonds are indicated by black dashed lines. The system mimics interactions between β-catenin and ICAT [PDB ID: 1LUJ]. *(C)* Interactions between beta-catenin (134-671) (gray cartoon) and compound 2 (bright orange sticks) or compound 3 (violet sticks). Interacting amino acids are indicated by gray lines; bonds are indicated by black dashes (PDB: 9I8X and 9I8W).

**Table 2.**
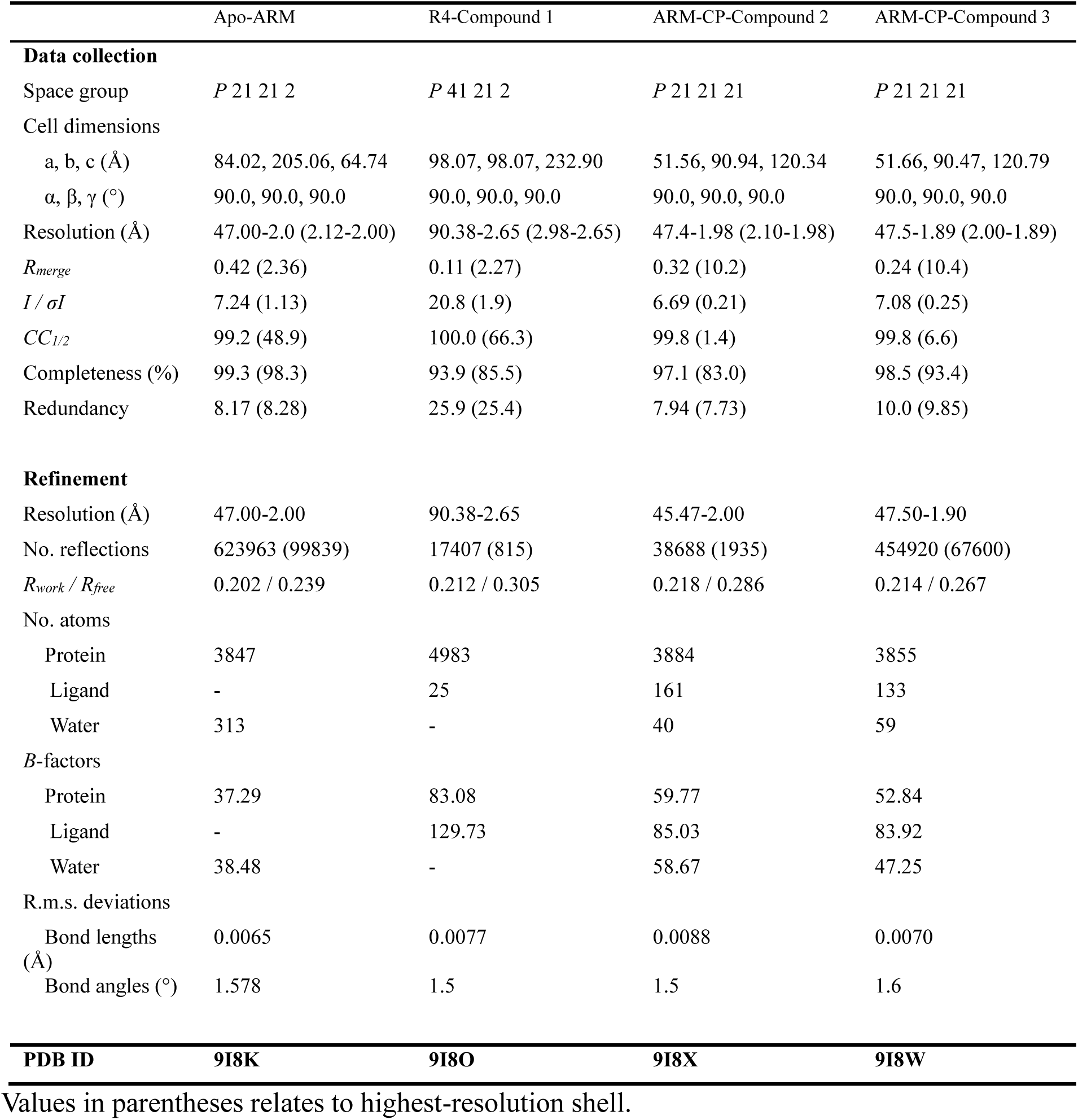
Data collection and refinement statistics for: apo beta-catenin (150-663) (Apo-ARM), beta-catenin R4 (141-305) in complex with Compound-1 (R4-Compound 1), beta-catenin (134-671) bound to cyclic peptide in complex with Compound 2 (ARM-CP-Compound 2) and beta-catenin (134-671) bound to cyclic peptide in complex with Compound 2 (ARM-CP-Compound 3).

|  | Apo-ARM | R4-Compound 1 | ARM-CP-Compound 2 | ARM-CP-Compound 3 |
| --- | --- | --- | --- | --- |
| **Data collection** |  |  |  |  |
| Space group | *P* 21 21 2 | *P* 41 21 2 | *P* 21 21 21 | *P* 21 21 21 |
| Cell dimensions |  |  |  |  |
| a, b, c (Å) | 84.02, 205.06, 64.74 | 98.07, 98.07, 232.90 | 51.56, 90.94, 120.34 | 51.66, 90.47, 120.79 |
| $\alpha$, $\beta$, $\gamma$ (°) | 90.0, 90.0, 90.0 | 90.0, 90.0, 90.0 | 90.0, 90.0, 90.0 | 90.0, 90.0, 90.0 |
| Resolution (Å) | 47.00-2.0 (2.12-2.00) | 90.38-2.65 (2.98-2.65) | 47.4-1.98 (2.10-1.98) | 47.5-1.89 (2.00-1.89) |
| *R*<sub>merge</sub> | 0.42 (2.36) | 0.11 (2.27) | 0.32 (10.2) | 0.24 (10.4) |
| *I* / σ*I* | 7.24 (1.13) | 20.8 (1.9) | 6.69 (0.21) | 7.08 (0.25) |
| *CC*<sub>1/2</sub> | 99.2 (48.9) | 100.0 (66.3) | 99.8 (1.4) | 99.8 (6.6) |
| Completeness (%) | 99.3 (98.3) | 93.9 (85.5) | 97.1 (83.0) | 98.5 (93.4) |
| Redundancy | 8.17 (8.28) | 25.9 (25.4) | 7.94 (7.73) | 10.0 (9.85) |
| **Refinement** |  |  |  |  |
| Resolution (Å) | 47.00-2.00 | 90.38-2.65 | 45.47-2.00 | 47.50-1.90 |
| No. reflections | 623963 (99839) | 17407 (815) | 38688 (1935) | 454920 (67600) |
| *R*<sub>work</sub> / *R*<sub>free</sub> | 0.202 / 0.239 | 0.212 / 0.305 | 0.218 / 0.286 | 0.214 / 0.267 |
| No. atoms |  |  |  |  |
| Protein | 3847 | 4983 | 3884 | 3855 |
| Ligand | - | 25 | 161 | 133 |
| Water | 313 | - | 40 | 59 |
| *B*-factors |  |  |  |  |
| Protein | 37.29 | 83.08 | 59.77 | 52.84 |
| Ligand | - | 129.73 | 85.03 | 83.92 |
| Water | 38.48 | - | 58.67 | 47.25 |
| R.m.s. deviations |  |  |  |  |
| Bond lengths (Å) | 0.0065 | 0.0077 | 0.0088 | 0.0070 |
| Bond angles (°) | 1.578 | 1.5 | 1.5 | 1.6 |
| **PDB ID** | **9I8K** | **9I8O** | **9I8X** | **9I8W** |

Values in parentheses relates to highest-resolution shell.

Although the cyclic peptide bound BC-ARM construct offers good quality crystals, resulting in high-resolution data, it does have a significant caveat as a ‘platform protein’ to be used in crystallography studies. The interactions between the peptide and the protein are occupying significant space of β-catenin surface that are involved in interactions with its physiological partners (*e.g.*, BCL9, E-cadherin, Tcf4, Axin), therefore restricting BC-ARM surface when performing crystallography fragment screening. Traditionally used apo construct of BC-ARM, that did not pose such limitations was reported to diffract in the range of 2.9-3.5 Å (PDB ID: 4HM9) (10), similarly in our hands this protein produced crystals of varying quality, routinely diffracting to 3 Å. This proved to be insufficient for high-throughput crystallography studies. Analysis of this construct led us to design and generation of the truncated version, to eliminate unstructured and flexible regions, with amino acids 50-663. Restricting BC-ARM motif ends allowed us to identify the crystallization conditions for the apo structure without the need for additional stabilizing partners. This construct can be standardly utilized to produce well diffracting crystals, suited for ligand co-crystallization or soaking in a reliable manner (**Figure 4**).

**Figure 4.**
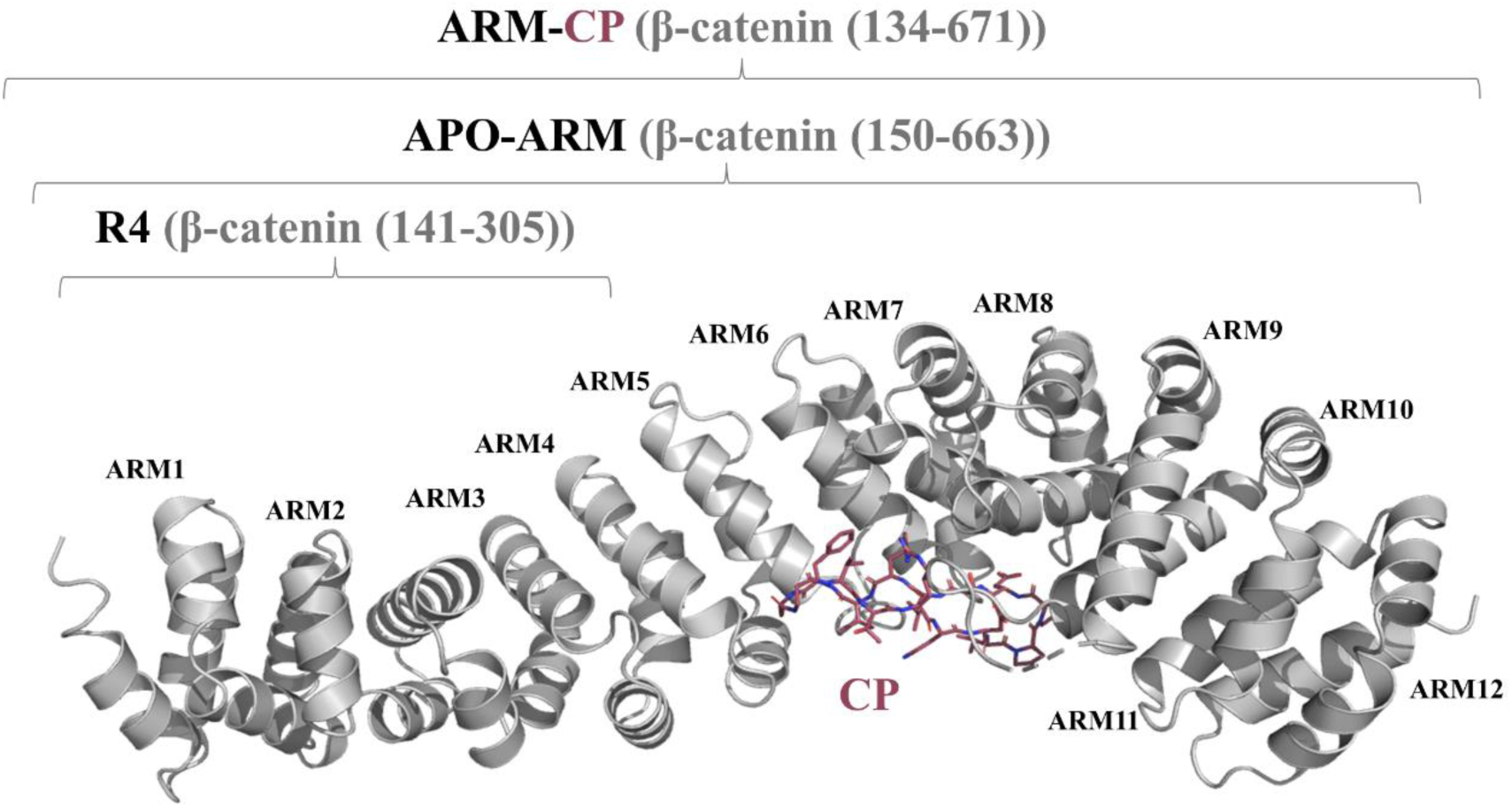
Selection of beta-catenin constructs used in this study. Figure was prepared based on ARM-CP structure which contains β-catenin (134-671) (ARM) protein shown as gray cartoon and cyclic peptide (CP) shown as raspberry sticks. ARM-CP crystallization system was utilized to obtain structures with compound 2 and compound 3. R4 (Beta-catenin (141-305)) construct containing first four armadillo repeats of β-catenin protein was utilized to obtain structure with compound 1. APO-ARM (Beta-catenin (150-663)) construct was designed based on previous experiments and utilized to obtain new apo structure (PDB: 9I8K).

In conclusion, we believe that the new ligand-bound β-catenin structures reported here, in conjunction with the novel β-catenin construct, establish a new platform for investigation of β-catenin. Furthermore, this new construct, when combined with the FragLites library or any other library, may represent an innovative approach for identifying novel hotspots for degradation of β-catenin.

## Material and methods

### BC-R4 (141–305) expression and purification

First four armadillo repeats (residues 141-305) of human β-catenin (CTNB1, UniProt: P35222) were cloned into a vector to generate a N-terminal 6xHis-Tag and SUMO domain [6xHis-SUMO-BC(R4)]. Vector was transformed into chemically competent recombinant *E. coli* BL21(DE3) cells and cultured in LB broth medium supplemented with 50 μg/mL kanamycin. Culture was incubated overnight at 37°C with aeration 160 rpm, till an optical density at 600 nm reached 0.5-0.6. Once the certain optical density was achieved the protein expression was induced by supplementing culture with isopropyl β-D-1-thiogalactopyranoside (IPTG) to a final concentration 1 mM. After expression induction culture was incubated overnight at 25°C with aeration 90 rpm before cells pellet harvesting by centrifugation at 6000×*g* for 20 minutes at 4°C. Cells were stored in −80°C until purification. To start purification cells pellet was resuspended in lysis buffer (50 mM Tris/HCl pH 8.0 at 4°C, 300 mM NaCl, 5% glycerol) supplemented with 10 mM β-ME (2-Mercaptoethanol, BME), 1 mM PMSF (Phenylmethanesulfonyl fluoride), 0.5 mM EDTA, cOmplete™ protease inhibitors EDTA-free cocktail tablet (Roche) and 2.5 U/mL PierceTM Universal Nuclease. Cells were disintegrated using EmulsiFlex. Afterwards polyethyleneimine was added to the lysate reaching 0.1% concentration. Lysate was clarified by centrifugation at 30,000 *g* for 30 minutes at 4°C. At the first step of purification Immobilized Metal Affinity Chromatography (IMAC) was performed. For this purpose, clarified lysate was supplemented with 5 mM imidazole before binding for 1 hour at 4°C with 10 ml Cytiva Ni Sepharose 6 Fast Flow resin equilibrated in 50 mM Tris/HCl pH 8.0 at 4°C, 300 mM NaCl, 5% glycerol, 5 mM imidazole and 10 mM BME. After binding resin was recovered from lysates using glass chromatography column and washed with 10 CV of 50 mM Tris/HCl pH 8.0 at 4°C, 300 mM NaCl, 5% glycerol, 5 mM imidazole and 10 mM BME. All fractions obtained during washing were collected. Elution was performed by suspension of the resin with bound protein in 2 CV of 50 mM Tris/HCl pH 8.0 at 4°C, 300 mM NaCl, 5% glycerol, 0.5 M imidazole and 10 mM BME and incubated for 5 minutes. Afterwards eluate was collected. Resin was additionally washed with 1 CV of 50 mM Tris/HCl pH 8.0 at 4°C, 300 mM NaCl, 5% glycerol, 0.5 M imidazole and 10 mM BME to collect the void volume and achieve higher recovery. Protein was cleaved overnight at 4℃ using 1:100 (Mass ratio protease:protein) Ulp1 protease with simultaneous dialysis against buffer consisting of 50 mM Tris pH 8.0 at 4℃, 300 mM NaCl and 10 mM BME. Additionally, second dialysis lasting 4 hours at 10℃ against 50 mM Tris pH 8.0 at 4℃, 300 mM NaCl, 10 mM BME was performed to decrease the imidazole concentration. To remove the fusion tag and the protease from the cleaved protein Anion Exchange Chromatography (AIEX) was performed. For this purpose, sample after second dialysis was diluted 10x in 50 mM Tris/HCl pH 8.0 at RT supplemented with 10 mM BME. Diluted sample was loaded into the Cytiva HiTrap Q HP 5 mL column equilibrated in 50 mM Tris/HCl pH 8.0 at 4°C, 30 mM NaCl and 10 mM BME and washed with 5 CV of 50 mM Tris/HCl pH 8.0 at 4°C, 30 mM NaCl and 10 mM BME. Then elution was performed using 50 mM Tris/HCl pH 8.0 at 4°C, 500 mM NaCl and 10 mM BME. For concentration additional IMAC was performed. Sample was loaded into the Cytiva HisTrap HP 5 mL column equilibrated in 50 mM Tris/HCl pH 8.0 at RT, 300 mM NaCl, 5% glycerol and 10 mM BME and washed with 5 CV of 50 mM Tris/HCl pH 8.0 at RT, 300 mM NaCl, 5% glycerol and 10 mM BME. Afterwards elution was performed with 14 CV of 50 mM Tris/HCl pH 8.0 at RT, 300 mM NaCl, 5% glycerol, 10 mM BME and imidazole gradient (0-150 mM). Then dialysis was performed twice to exchange the buffer for protein storage – first dialysis was incubated overnight while second for 4 hours, both at 10℃. Each dialysis was performed against buffer consisting of 50 mM Tris pH 8.0 at 4℃, 300 mM NaCl and 10 mM BME. Size exclusion chromatography was performed as the final purification step. For this purpose, sample after dialysis was loaded on the Cytiva HiLoad 26/600 Superdex 75 pg column equilibrated in 20 mM Tris/HCl pH 8.0 at 4°C, 150 mM NaCl and 1 mM TCEP. Eluted fractions containing protein were flash frozen in liquid nitrogen. Samples were stored at −80°C until further use.

### BC-ARM (134–671) and shortened BC-ARM (150–663) expression and purification

Armadillo domain (residues 134-671) or shortened armadillo domain (residues 150-663) of human β-catenin (CTNB1, UniProt: P35222) were cloned into the vector to generate a N-terminal 6xHis-Tag and SUMO domain [6xHis-SUMO-BC(ARM)]. Moreover, for armadillo domain (residues 134-671) of human β-catenin a version containing AviTag [6xHis-SUMO-BC(ARM)-AviTag] was generated for further *in vivo* biotinylation and biophysical analysis (immobilization in SPR experiments). Vectors were transformed into a chemically competent recombinant *E. coli* BL21(DE3) cells and cultured in LB broth medium supplemented with 50 μg/mL kanamycin. For BC-ARM version with AviTag additionally biotinylation was performed by co-expression of the protein of interest with *E. coli* biotin ligase BirA using a single vector and biotin supplementation of the culture medium. Cultures were incubated overnight at 37°C with aeration 160 rpm, till an optical density at 600 nm reached 0.5-0.6. Once required optical density was achieved protein expression was induced by supplementing cultures with isopropyl β-D-1-thiogalactopyranoside (IPTG) to a final concentration 0.5 mM. After expression induction cultures were incubated overnight at 25°C with aeration 90 rpm before cells pellet harvesting by centrifugation at 6000×*g* for 20 minutes at 4°C. Cells were stored in −80°C until purification. To start purification cells pellets were resuspended in lysis buffer (for BC-ARM protein - 50 mM Tris/HCl pH 8.0 at 4°C, 300 mM NaCl, 10% glycerol while for shortened BC-ARM protein - 20 mM potassium phosphate pH 8.0 at 4°C, 300 mM NaCl, 10% glycerol, 20mM imidazole) supplemented with β-ME (2-Mercaptoethanol; 10 mM for BC-ARM while 5 mM for shortened BC-ARM), 1 mM PMSF (Phenylmethanesulfonyl fluoride), 0.5 mM EDTA, cOmplete™ protease inhibitors EDTA-free cocktail tablet (Roche) and 5 U/mL PierceTM Universal Nuclease. Cells were disintegrated using EmulsiFlex. Afterwards polyethyleneimine was added to lysates reaching 0.1% concentration. Lysates were clarified by centrifugation at 30,000 *g* for 30 minutes at 4°C. At the first step of purification Immobilized Metal Affinity Chromatography was performed. For this purpose, clarified lysates were supplemented with 5 mM imidazole before binding for 1 hour at 4°C with 10 ml Cytiva Ni Sepharose 6 Fast Flow resin equilibrated in 50 mM Tris/HCl pH 8.0 at 4°C, 300 mM NaCl, 10% glycerol, 5 mM imidazole and 10 mM BME. After binding resin was recovered from lysates using glass chromatography column and washed with 10 CV of 50 mM Tris/HCl pH 8.0 at 4°C, 300 mM NaCl, 5% glycerol, 20 mM imidazole and 10 mM BME. Elution was performed by suspension of the resin with bound proteins in 2 CV of 50 mM Tris/HCl pH 8.0 at 4°C, 300 mM NaCl, 5% glycerol, 0.3 M imidazole and 10 mM BME and incubated for 5 minutes. Afterwards eluates were collected. Resin was additionally washed with 1 CV of 50 mM Tris/HCl pH 8.0 at 4°C, 300 mM NaCl, 5% glycerol, 0.3 M imidazole and 10 mM BME to collect the void volume and achieve higher recovery. Proteins were cleaved overnight at 4℃ using 1:100 (Mass ratio protease:protein) Ulp1 protease with simultaneous dialysis against buffer consisting of 20 mM Sodium Phosphate pH 8.0 at 4℃, 300 mM NaCl, 2 mM DTT. Size Exclusion Chromatography was performed as the final purification step. For this purpose, sample containing BC-ARM after dialysis was loaded into the Cytiva HiLoad 26/600 Superdex 75 pg column equilibrated in 25 mM sodium phosphate pH 7.4 at RT, 300 mM NaCl, 2.5 % glycerol and 5 mM DTT while sample containing shortened BC-ARM after dialysis was loaded into the HiLoad 26/600 Superdex 200 pg column equilibrated in 25 mM sodium phosphate pH 8.0 at RT, 300 mM NaCl and 2 mM DTT. For crystallography purposes protein was loaded into a column equilibrated in 20 mM Tris/HCl pH 8.0 at 4°C, 150 mM NaCl and 1 mM TCEP. Eluted fractions containing protein were flash frozen in liquid nitrogen. Samples were stored at −80 °C until further use.

### Crystallization

Crystallization trials have been conducted at 19°C applying hanging-drop vapor diffusion method by mixing protein with an equal volume of reservoir solution. Crystals of apo beta-catenin (150–663) (Apo-ARM) at 9.5 mg/ml formed after 2 days in Morpheus buffer system 1 pH 6.5, 0.12 M monosaccharides mix, 50% precipitant mix 4; crystals of beta-catenin R4 (141–305) at 8 mg/ml in complex with Compound-1 (R4-Compound 1) formed within 3 days in 0.14 M ammonium acetate, 0.7 M MES pH 6.5, 21 % glycerol ethoxylate; crystals of beta-catenin (134-671) at 5 mg/ml bound to cyclic peptide (1:1.4 molar excess) in complex with Compound 2 (ARM-CP-Compound 2) formed within 3 days in 0.1 M Bis-TRIS pH 6.5, 0.2 M lithium sulfate monohydrate, 26% PEG 3350; and crystals of beta-catenin (134-671) at 5 mg/ml bound to cyclic peptide (1:1.4 molar excess) in complex with Compound 3 (ARM-CP-Compound 3) formed within 3 days in 0.1 M ADA pH 6.5, 100 mM ammonium sulfate, 30% PEG MME 5000. Ligands were added to protein in 1:1.3 molar excess and incubated for 1 h at room temperature. All crystals were flash frozen with the addition of 20% (v/v) ethylene glycol.

### Structure determination

Data for apo beta-catenin (150-663) (Apo-ARM), beta-catenin R4 (141-305) in complex with Compound-1 (R4-Compound 1), beta-catenin (134-671) bound to cyclic peptide in complex with Compound 2 (ARM-CP-Compound 2) and beta-catenin (134-671) bound to cyclic peptide in complex with Compound 3 (ARM-CP-Compound 3) were collected at the Deutsches Elektronen-Synchrotron DESY (Hamburg, Germany) (21,22) to a resolution of 1.9 – 2.65 Å and processed automatically utilizing XDS package at the beamline (23). Initial phases were obtained by molecular replacement with PHASER (24) using Beta-catenin armadillo repeat domain bound to ICAT structure (PDB ID: 1M1E (25)) as a search model. The model was built using COOT (26) and refined using REFMAC5 (27). Ligand topologies were generated with *AceDRG* (*28*). All figure models were generated using PyMOL (29).

### Surface Plasmon Resonance (SPR)

The molecular interactions were tested by Surface Plasmon Resonance (SPR) using Biacore 8K instrument (Cytiva). The experiments were performed on the CM5 series S Biosensor chip (Cytiva) in all channels (channel 1-8). To start with the immobilization of NeutrAvidin was performed on the CM5 chip using standard amine coupling kit (Cytiva) with a flow rate of 10 µL/min chip (Cytiva) using standard capture protocol. After the capture, the surface was equilibrated for 2-4 hours in the running buffer (PBS pH 7.4, 0.05% Tween-20, 1mM DTT, 2% DMSO). Finally, the surface of flow cell 2 was conditioned by three injections of regeneration solution (1M NaCl in 50 mM NaOH). In the next step a solution of a biotinylated BC-ARM (amino acid region 134-671) at concentration of 80 µg/ml in the immobilization buffer (PBS pH 7.4, 0.05% Tween20, 1mM DTT) was injected over the sensor surface for 180 s. The immobilization levels were assessed and oscillated at about 5000-6000 RU in each channel. Subsequently, both flow cells were blocked with 50 µM biotin-PEO3 (Biotium) for 90 s at 10 µL/min. All parts of the flow system except for the sensor chip were washed with 50% isopropanol in 1M NaCl and 50 mM NaOH. For the dose-response experiments, the stock solutions of either 20 mM or 100 mM compounds were loaded onto a 384-well LDV plate (Labcyte Inc.). Compounds were transferred to the 384-well SPR destination plate (781280, Greiner) by the Echo® 555 liquid handler (Labcyte Inc.) to create dose-response curves. For fragments, the curves consisted of 10 points with concentrations ranging from 50 to 1000 µM (50, 70, 100, 135, 190, 265, 370, 515, 720, 1000 µM) and for other compounds, the curves consisted of 10 points with concentrations ranging from 1 to 100 µM (1, 2, 5, 10, 17.5, 25, 37.5, 50, 75, 100 µM). Each well of the destination plate, containing 1010 nL of the compounds backfilled with DMSO, was topped up with 99 µL of sample buffer (PBS pH 7.4, 0.05% P-20, 1 mM DTT, 1% DMSO) using a MultiFlo FX Dispenser (BioTek). The prepared plates were then centrifuged at 1000 rcf for 1 min at room temperature using the 5804R centrifuge (Eppendorf). A dedicated analysis method was configured in the Biacore 8K Control Software (Cytiva), which included an association time of 60s, a dissociation time of 120 s, and a flow rate of 30 µL/min. The method also incorporated positive, negative and a carry-over controls, and solvent correction consisting of 10 points. The results were analyzed using the dedicated method in Biacore Insight Evaluation Software (Cytiva). In multi-cycle kinetics experiments, steady-state affinity for fragments was determined based on analyte binding early stability point (15 seconds from the beginning of injection), while for other compounds, it was determined based on analyte binding late stability point (5 seconds before the end of injection). In single-cycle kinetics experiments, the equilibrium dissociation constant was determined by using a 1:1 binding model.

### Differential Scanning Fluorimetry (DSF)

The effect of the compounds on stabilization of the BC-ARM (amino acid region 134-671) was investigated with the Differential Scanning Fluorimetry (DSF) assay. First, the BC-ARM protein was thawed and centrifuged at 18,000 rcf at 4°C for 5 min (Centrifuge 5430R, Eppendorf). The protein concentration in the supernatant was measured (BioSpectrometer Basic, Eppendorf) and used to prepare a master mix containing 5 μM of BC-ARM in the assay buffer PBS, pH=7.4, 1 mM DTT. The master mix solution was pipetted into a 384-well PCR plate (4titude®, FrameStar®). The plates were spun down at 1000 rcf for 10 sec at room temperature (Centrifuge 5804R, Eppendorf). Subsequently, SYPRO™ Orange Protein Gel Stain (cat. nr S6650, Thermo Scientific) and tested compounds were added to the plate using Echo® 555 liquid handler (Labcyte Inc.). Fragments were tested at the concentration of 500 μM and other compounds at 100 μM in the final reaction mixture of 10 μL per well. SYPRO™ Orange was used at the final dilution of 7.5x, and the DMSO content was adjusted to 5%. The plate was spun down at 1000 rcf at 22°C for 1 min (Centrifuge 5804R, Eppendorf), shaken by Vibroturbulator (Union Scientific) at level 10 for 1 min, spun down again, and incubated in the darkness for 15 min at room temperature. The readout was done using ViiA™ 7 Real-Time PCR System (Applied Biosystems) with a ramp rate of 0.1°C/s and the temperature range from 25°C to 95°C. The ΔTm and SSMD values were determined according to published procedures (SSMD – (30); dTM – (31).

### Chemical synthesis

**Scheme 1.**
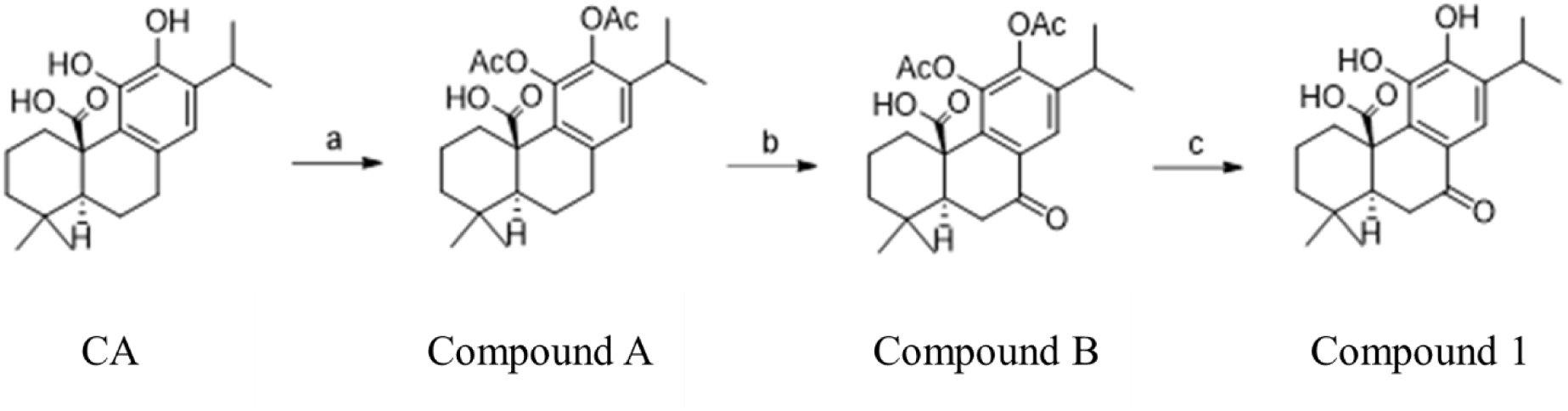
Synthesis of Compound 1^a^. ^a^ Reagents and conditions: (a) Ac_2_O, Pyridine, rt, 48 h; (b) CrO_3_, AcOH, rt, 18 h; (c) MeNH_2_, EtOH, rt, 4 h.

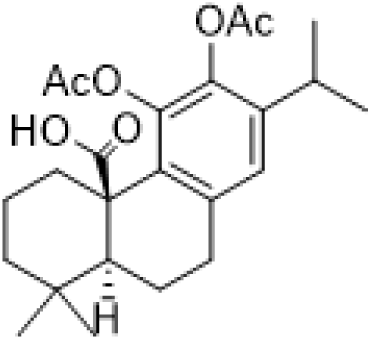
(4aR,10aS)-5,6-diacetoxy-7-isopropyl-1,1-dimethyl-1,3,4,9,10,10a-hexahydrophenanthrene-4a(2H)-carboxylic acid (Compound A). Carnosic acid (3.00 g, 9.02 mmol, 1.00 eq) was dissolved in pyridine (21.00 mL). Acetic anhydride (2.5 mL, 26.49 mmol, 3.00 eq) was slowly added into the solution at room temperature, then the reaction mixture was stirred at room temperature for 24 h under Ar atmosphere. After this time more acetic anhydride (4.9 mL, 51.95 mmol, 6.00 eq) was added to ensure full conversion and the mixture was stirred for another 24 h. After completion, the mixture was diluted with 10% aqueous citric acid solution and extracted with EtOAc. Organic fractions were combined, washed with water and brine, dried over anhydrous Na_2_SO_4_, filtered, and concentrated under reduced pressure. The crude product was purified by silica gel flash chromatography (gradient from 0% to 20% of EtOAc in hexanes) to give pure C**ompound A** (3.06 g, 7.35 mmol, 85%) as a pale yellow powder.

^1^H NMR (500 MHz, DMSO) δ 12.31 (s, 1H), 6.99 (s, 1H), 3.24 – 3.05 (m, 1H), 2.91 – 2.77 (m, 3H), 2.35 – 2.27 (m, 1H), 2.25 (s, 3H), 2.18 (s, 3H), 2.08 – 1.97 (m, 1H), 1.79 – 1.72 (m, 1H), 1.51 – 1.39 (m, 3H), 1.28 – 1.20 (m, 1H), 1.14 (d, *J* = 6.9 Hz, 3H), 1.08 (d, *J* = 6.9 Hz, 3H), 1.06 – 1.01 (m, 1H), 0.94 (s, 3H), 0.81 (s, 3H). MS m/z [M-H]^-^ 415.3

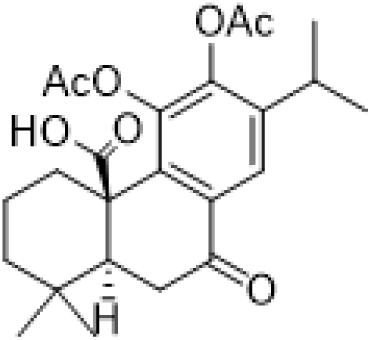
(4aR,10aS)-5,6-diacetoxy-7-isopropyl-1,1-dimethyl-9-oxo-1,3,4,9,10,10a-hexahydrophenanthrene-4a(2H)-carboxylic acid (Compound B). (4aR,10aS)-5,6-diacetoxy-7-isopropyl-1,1-dimethyl-1,3,4,9,10,10a-hexahydrophenanthrene-4a(2H)-carboxylic acid (0.32 g, 0.69 mmol, 1.00 eq) was dissolved in acetic acid (6.90 mL), then chromium(VI) oxide (0.21 g, 2.07 mmol, 3.00 eq) was added in one portion and the reaction mixture was stirred at room temperature for 18 h, after which time full conversion was observed. Reaction mixture was diluted with EtOAc, washed with water and brine, dried over anhydrous Na_2_SO_4_, filtered and concentrated under reduced pressure. The crude product was purified by silica gel flash chromatography (gradient from 0% to 15% of EtOAc in hexanes) to give pure **Compound B** (0.22 g, 0.46 mmol, 67%) as a pale yellow solid.

^1^H NMR (700 MHz, CDCl_3_) δ 8.05 (s, 1H), 3.39 – 3.34 (m, 1H), 3.31 (dd, *J* = 17.2, 15.2 Hz, 1H), 2.94 (hept, *J* = 6.9 Hz, 1H), 2.70 (dd, *J* = 17.2, 3.3 Hz, 1H), 2.29 (s, 3H), 2.26 (s, 3H), 2.18 – 2.14 (m, 1H), 2.14 – 2.06 (m, 1H), 1.63 – 1.57 (m, 1H), 1.53 – 1.48 (m, 1H), 1.33 (tdd, *J* = 13.3, 8.4, 4.3 Hz, 2H), 1.25 (d, *J* = 6.9 Hz, 3H), 1.17 (d, *J* = 7.0 Hz, 3H), 0.95 (s, 3H), 0.90 (s, 3H). MS m/z [M+H]^+^ 431.2

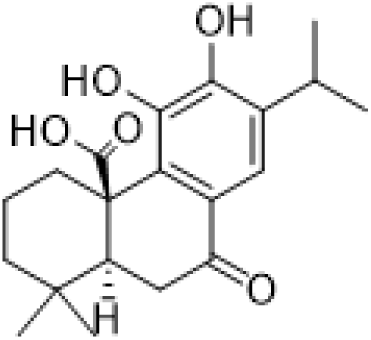
(4aR,10aS)-5,6-dihydroxy-7-isopropyl-1,1-dimethyl-9-oxo-1,3,4,9,10,10a-hexahydrophenanthrene-4a(2H)-carboxylic acid (Compound 1). (4aR,10aS)-5,6-diacetoxy-7-isopropyl-1,1-dimethyl-9-oxo-1,3,4,9,10,10a-hexahydrophenanthrene-4a(2H)-carboxylic acid (0.39 g, 0.88 mmol, 1.00 eq) was dissolved in EtOH (5.00 mL), then a mixture of methylamine in EtOH (1.1 mL 33% wt in EtOH, ∼8.80 mmol, ∼10.00 eq) was added to the reaction mixture over 10 min and the reaction was vigorously stirred for 4 h at room temperature. After this time, the mixture was concentrated under reduced pressure and the crude mixture was purified by silica gel flash chromatography (gradient from 20% to 30% of EtOAc in hexanes) to give pure **Compound 1** (0.15 g, 0.43 mmol, 50%) as a yellow powder.

^1^H NMR (400 MHz, CDCl_3_) δ 7.73 (s, 1H), 6.33 (br, 2H), 3.44 (d, *J* = 13.9 Hz, 1H), 3.41 – 3.31 (m, 1H), 3.27 – 3.18 (m, 1H), 2.67 (dd, *J* = 17.2, 3.3 Hz, 1H), 2.14 (dd, *J* = 15.2, 3.4 Hz, 1H), 1.88 – 1.72 (m, 1H), 1.72 – 1.60 (m, 1H), 1.59 – 1.47 (m, 1H), 1.43 – 1.31 (m, 1H), 1.31 – 1.25 (m, 1H), 1.22 (d, *J* = 3.2 Hz, 3H), 1.21 (d, *J* = 3.2 Hz, 3H), 0.97 (br, 6H). MS m/z [M+H]^+^ 347.20, [M-H]^-^ 345.10

## Accession codes

All atomic coordinates and crystallographic results of β-catenin/small molecule complexes and for new structure of APO-β-catenin were deposited to Protein Data Bank as: 9I8O, 9I8X, 9I8W and 9I8K respectively; and are accessible from their websites (*i.e*. https://www.rcsb.org/).

## Acknowledgements

We acknowledge DESY (Hamburg, Germany), a member of the Helmholtz Association HGF, for the provision of experimental facilities. Parts of this research were carried out at PETRA III, and we would like to thank Dr. Eva Crosas, Dr Johanna Hakanpaa and Dr. Spyridon Chatziefthymiou for their assistance in using beamline P11. Beamtime was allocated for proposal P-20010353. This research was funded in part by The National Centre for Research and Development, grant number POIR.01.02.00-00-0073/18. The authors would like to extend their gratitude to Tomas Drmota and Przemyslaw Glaza for the proofreading and valuable suggestions.

## Authors contributions

Conceptualization of the manuscript and writing: MK, ANS, MWP; Review and edition of the manuscript: MJW; Chemical synthesis: JS, FU; Protein purification: ANS, GD, DG, MK, MS, IHW, JW; Crystallography and structure determination: MK, ANS, JW, KMGM; Biophysical assays: DK; Research supervised by: MB, SC, MJW.

## Assistive writing technologies

We used assistive writing technologies of Google Gemini during manuscript preparation. This supplementary tool acted as an editor, not as drivers of content creation. The listed authors thoroughly reviewed, revised, and selectively implemented suggested edits to ensure accuracy, consistency, and clarity. The responsibility for the content and quality of the manuscript remains with the authors.

## Abbreviations

APC: Adenomatous Polyposis Coli
BC: beta catenin
BCL9: B-cell lymphoma 9
CBP: CREB Binding Protein
*E. coli*: *Escherichia coli*
DSF: Differential Scanning Fluorimetry
DMSO: Dimethyl Sulfoxide
DTT: Dithiothreitol
EDTA: Ethylenediaminetetraacetic Acid
SBDD: Structure-Based Drug Design
SPR: Surface Plasmon Resonance
TPD: Targeted Protein Degradation
TCF/LEF family: T-cell factor/lymphoid enhancer factor
UPS: Ubiquitin-Proteasome System
PDB: Protein Data Bank
Wnt: combination of “Wingless” (a Drosophila gene) and “Integration site” (referring to Int-1, a gene in mice)

